# YOLO-Behaviour: A simple, flexible framework to automatically quantify animal behaviours from videos

**DOI:** 10.1101/2024.08.26.609387

**Authors:** Alex Hoi Hang Chan, Prasetia Putra, Harald Schupp, Johanna Köchling, Jana Straßheim, Britta Renner, Julia Schroeder, William D. Pearse, Shinichi Nakagawa, Terry Burke, Michael Griesser, Andrea Meltzer, Saverio Lubrano, Fumihiro Kano

## Abstract

Manually coding behaviours from videos is essential to study animal behaviour but it is labour-intensive and susceptible to inter-rater bias and reliability issues. Recent developments of computer vision tools enable the automatic quantification of behaviours, supplementing or even replacing manual annotations. However, widespread adoption of these methods is still limited, due to the lack of annotated training datasets and domain-specific knowledge required to optimize these models for animal research. Here, we present YOLO-Behaviour, a flexible framework for identifying visually distinct behaviours from video recordings. The framework is robust, easy to implement, and requires minimal manual annotations as training data. We demonstrate the flexibility of the framework with case studies for event-wise detection in house sparrow nestling provisioning, Siberian jay feeding, human eating behaviours, and frame-wise detections of various behaviours in pigeons, zebras, and giraffes. Our results show that the framework reliably detects behaviours accurately, and retrieve comparable accuracy metrics to manual annotation. However, metrics extracted for event-wise detection were less correlated with manual annotation, and potential reasons for the discrepancy between manual annotation and automatic detection are discussed. To mitigate this problem, the framework can be used as a hybrid approach of first detecting events using the pipeline and then manually confirming the detections, saving annotation time. We provide detailed documentation and guidelines on how to implement the YOLO-Behaviour framework, for researchers to readily train and deploy new models on their own study systems. We anticipate the framework can be another step towards lowering the barrier of entry for applying computer vision methods in animal behaviour.

## Introduction

Ever since the popularization of video cameras, animal researchers have been using videos to capture the behaviours of animals in captivity and in the field. Further propelled by user friendly video annotation tools like BORIS (Friard and Gamba, 2016), taking videos of animals and subsequently annotating for specific behaviours have become an essential part of data collection pipelines in animal behaviour. However this approach is time consuming (Chan et al., 2024) and can be susceptible to low observer reliability and repeatability (Tuyttens et al., 2014). To solve these problems, advances in computer science have leveraged large video datasets to create computer vision-based solutions to automate the quantification of behaviours from animal videos, leading to a significant shift in the scale and efficiency of extracting behaviours from video data (Couzin and Heins, 2022; Mathis and Mathis, 2020).

There are a few general approaches for automatically quantifying animal behaviours from videos. The first approach starts with 2D or 3D keypoint estimation on animals in a video frame, then use supervised (Wittek et al., 2022) or unsupervised (Graving and Couzin, 2020; Hsu and Yttri, 2021) methods to quantify behaviours using predicted keypoint information. Keypoint estimation of animal body parts from videos has recently been popularized with the development of tools including DeepLabCut (Mathis et al., 2018), SLEAP (Pereira et al., 2022) and DeepPoseKit (Graving et al., 2019), allowing fine-scaled body postures of animals to be measured precisely. However, keypoint estimation methods are often limited to captive settings (but see Chimento et al., 2024; Joska et al., 2021; Waldmann et al., 2024), and obtaining large keypoint ground truth datasets is often labour intensive.

The second approach is to directly input video frames into neural networks, and output observed behaviours. With recent benchmark datasets like animal kingdom (Ng et al., 2022), KABR (Kholiavchenko et al., 2024), PanAf20k (Brookes et al., 2024) or MammalNet (Chen et al., 2023), there is a growing trend of directly using video input for behavioural classification, even though the accuracy of such methods is often low (e.g., 50-60%; Kholiavchenko et al., (2024) table 1). Finally, a method for behaviour classification in more visually noisy scenes is to first isolate an animal in the video frame with a bounding box or mask, then input the cropped animal into a neural network classifier to classify behaviours (Lei et al., 2022; Yang et al., 2019). While this method is promising, with developed tools like LabGym (Goss et al., 2024; Hu et al., 2023), such approaches require among others segmentation masks and behavioural annotations of sequences as training data, which can be laborious to collect.

**Table 1:**
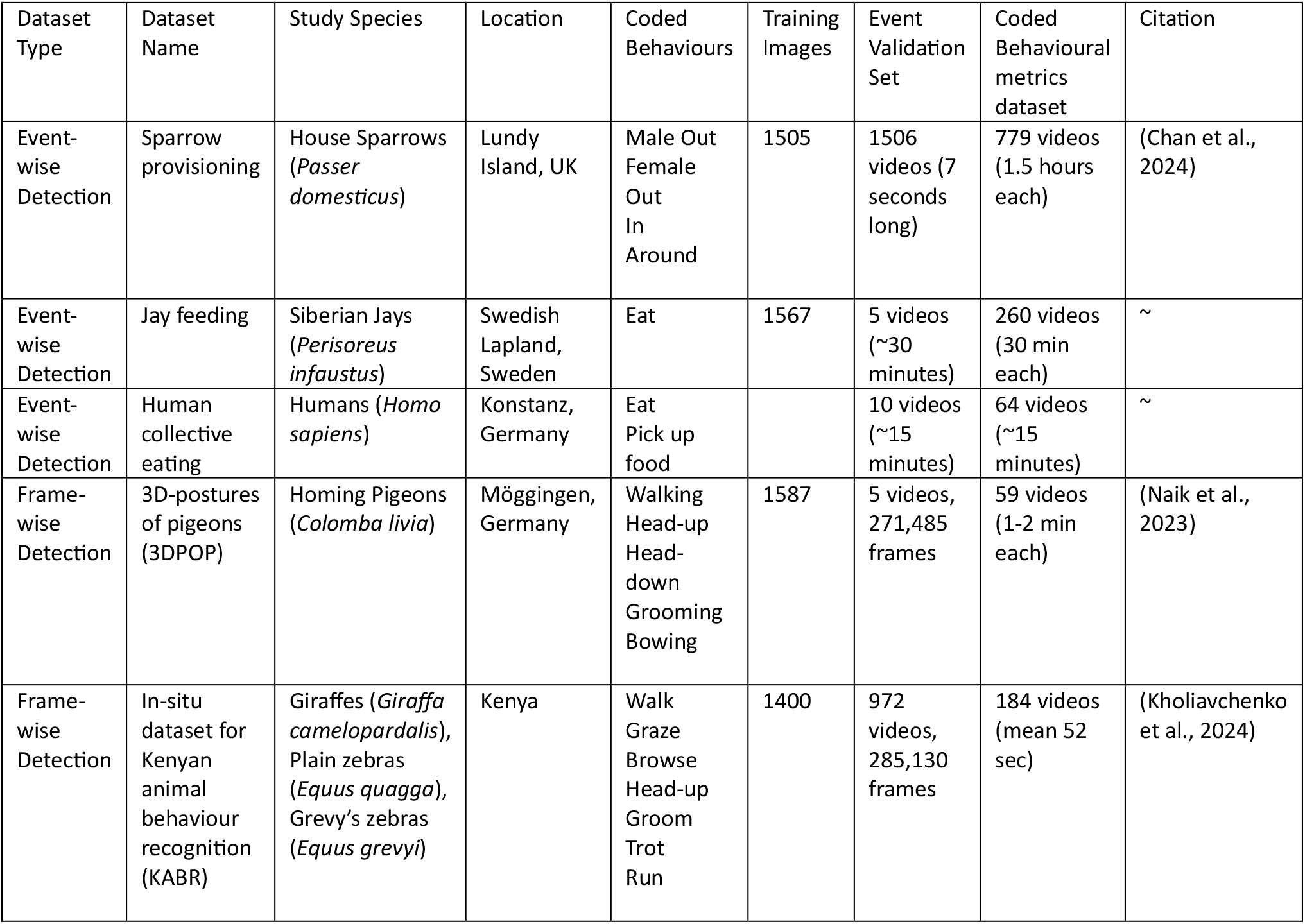
Details of datasets used in the current study.

Computer vision methods have been shown to be useful for quantifying behaviours in different species, but there is a lack of agreement on the most effective method for any given dataset and study system. Yet, especially under a rapidly changing climate and biodiversity crisis, it is more important than ever to leverage developments in computer vision to aid data collection on fundamental behavioural monitoring (Christin et al., 2019; Tuia et al., 2022), including individual level behaviours such as feeding rates, visit rates, or activity budgets, up to population level metrics. Such advances are not only important for deepening our understanding of biological systems, but to also gain insight into species conservation, and importantly increase the efficiency and wealth of data that can be collected and processed (Dell et al., 2014; Weinstein, 2018). However, while computer vision tools can be highly capable, they are often effective only in very specific contexts. These contexts are defined by unique characteristics such as the experimental setup, camera angles, lighting conditions, subject size and occlusions, all of which influence the suitability of a given method. Moreover, most frameworks require large amount of effort to collect training data, and sophisticated workflows to achieve automated behavioural coding (e.g., first training a keypoint model, then fitting an unsupervised algorithm, followed by training a supervised classifier; see Hsu and Yttri, 2021). Consequently, the domain-specific nature of current open-source computer vision algorithms poses a significant barrier to their widespread adoption by biologists and psychologists.

To overcome these limitations, we present the YOLO-Behaviour framework, an automatic behavioural detection and classification tool based on the common object-detector YOLOv8 (Jocher et al., 2023). An object detector is a class of models in computer vision that aims to localize an object within an image, by predicting a bounding box around a given object and its class. We leverage this model type to detect visually distinct behaviours in static frames, by providing training data of bounding boxes around behaviours within an image. The simplicity and robustness of the framework allows for widespread training and deployment by biologists with minimal training data and coding expertise, as demonstrated by the 5 case studies across study systems and taxa, which includes event-wise detection in 1) house sparrow (*Passer domesticus)* nestling provisioning, 2) Siberian jay (*Perisoreus infaustus)* feeding, 3) humans (*Homos sapiens)* eating; and frame-wise detections in 4) foraging pigeons (*Colomba livia)* as well as 5) roaming zebras (*Equus quagga* and *E. grevyi)* and giraffes (*Giraffa camelopardalis)*. Furthermore, we provide detailed code and documentation to facilitate its implementation in new study systems. We hope the simplicity of the proposed pipeline can promote the adoption of computer vision in animal researchers, thereby reducing the time required for manual coding in behavioural research and species monitoring.

## Materials and Methods

Below, we first describe the datasets we used to evaluate the YOLO-Behaviour framework, and the ways we categorized the datasets for evaluation. Next, we describe the whole pipeline, from training to post-processing and to optimization. Finally, we introduce the methods for evaluating the pipeline in terms of its ability to detect events accurately and to retrieve coded behavioural metrics.

### Datasets

We tested the robustness and generalization ability of YOLO-Behaviour by applying the method across five study systems across various taxa (Figure 1). Table 1 shows details of each dataset used, behaviours coded and size of training validation and test sets. We refer to the supplementary methods for detailed justification and description of each study system and data manipulation procedures. For each case study, we defined three dataset types, which differ slightly from conventional data splitting procedures. 1) Training images: annotated images used for training the YOLOv8 models. 2) Event validation set, a small number of videos for optimizing hyper-parameters and detailed evaluation of detection accuracy. 3) Coded behavioural metrics dataset, the largest dataset available to evaluate how the method can estimate coded behavioural metrics (e.g., feeding or visit rate), when compared with human annotation. All annotations from the training set are publicly available, except for the human dataset, which will not be available due to privacy and ethical concerns.

**Figure 1:**
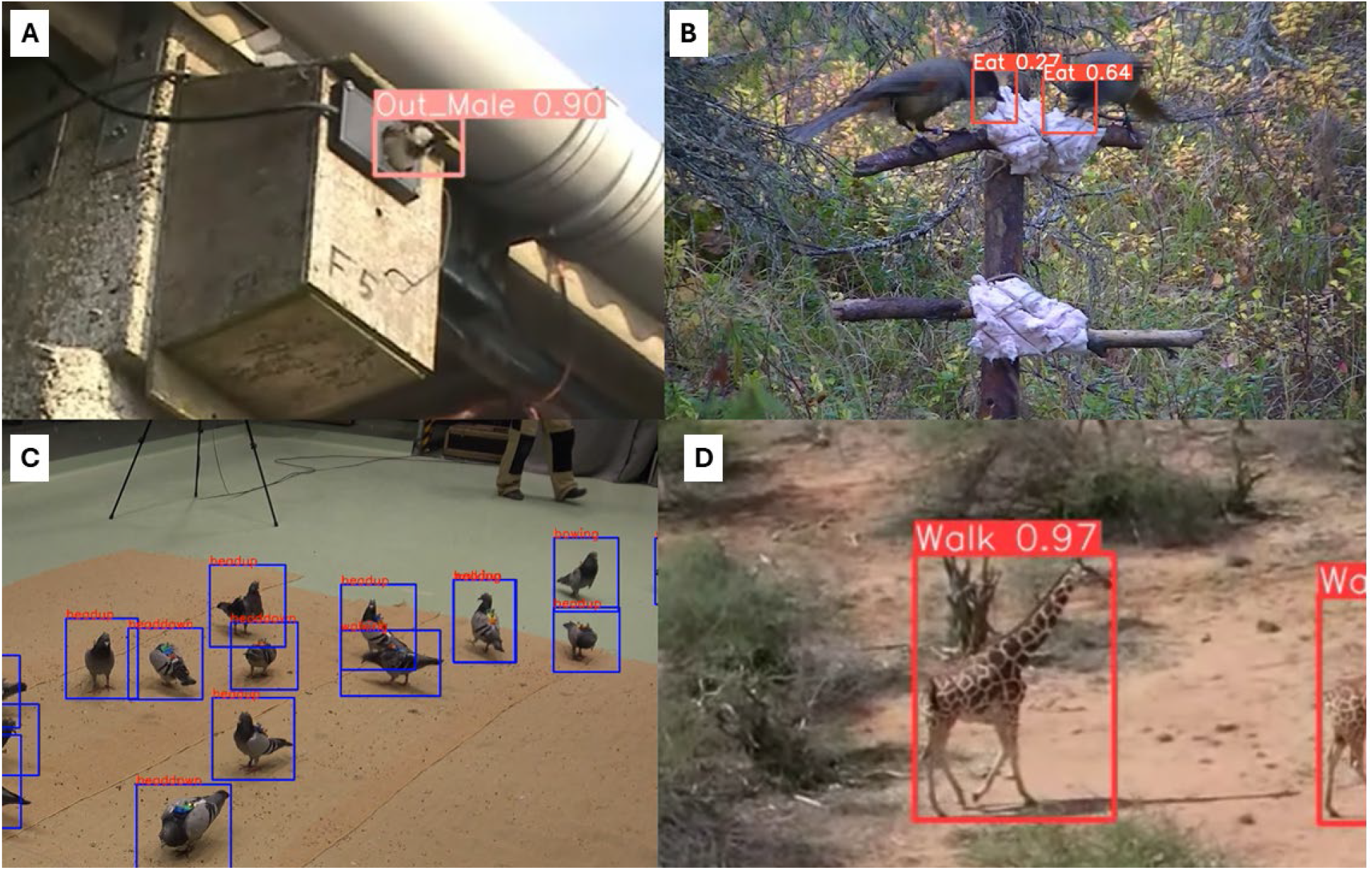
Case studies used to test the YOLO-Behaviour framework. Predictions of the YOLO model is overlayed onto each case study, with the predicted bounding box, class and model confidence. Human case study not shown due to privacy issues. A) House sparrow (*Passer domesticus*) provisioning videos collected on Lundy Island, UK. B) Siberian jay (*Perisoreus infaustus*) feeding videos collected in Swedish Lapland C) Homing pigeons (*Colomba livia*) behaviours collected in Möggingen, Germany, based on the 3D-POP dataset. D) Giraffes (*Giraffa camelopardalis*) behaviours collected in Mpala research center, Kenya, part of the KABR dataset.

### YOLO-Behaviour

The complete YOLO-Behaviour framework can be separated into three parts. 1) Data annotation and training: images were annotated and a YOLOv8 model was trained. 2) Post-processing: YOLO detections were processed and grouped using a tracking algorithm and 3) Optimization: a grid-search algorithm was used on the validation dataset to determine the best hyper-parameters for the final pipeline. We describe each of the steps in detail below. All code and documentation to apply the framework to novel systems can be found in the following link: https://alexhang212.github.io/YOLO_Behaviour_Repo/

#### 1) Data Annotation and Training

Firstly, random frames were sampled from each dataset, and bounding boxes were manually annotated for behaviours of interest, ensuring the bounding box encloses a visually distinctive part of the image that characterizes the behaviour. For example, in the Siberian jay and human eating datasets, the eating behaviour was captured by annotating a bounding box around the hand/ beak touching the mouth/ food respectively (Figure 1BC). For the pigeon and zebra/giraffe datasets, no manual annotation was done, since bounding boxes and behavioural labels were extracted using the dataset provided. We refer to supplementary methods for a detailed description of each dataset. After frames were extracted and annotated, the data was further split into training, validation and test sets using a 70%, 20%, 10% split. A YOLOv8-large model was then trained using the Ultralytics python package (Jocher et al., 2023), with default augmentation and training parameters.

#### 2) Post-processing

Once the YOLO models were trained, the models were used to detect behaviours from videos. However, the raw output of YOLO is bounding boxes with a given label and position in the frame, which is uninformative and can represent multiple behaviours being detected on the same frame (e.g., two jays pecking at the food from both sides). To solve this problem, we used the tracking algorithm SORT (Bewley et al., 2016) to group bounding box detections across spatial and temporal scales, by connecting closely detected bounding boxes. SORT is a widely used multi-object tracking algorithm, which is traditionally used for tracking the trajectory of detected objects in a video (e.g., human pedestrians walking in a video). Here, instead of tracking objects across the screen, we make use of the same type of algorithm to group bounding boxes of the same class to be classified as a single behavioural event. While other tracking algorithms exist with different complexity, we decided to use SORT due to its fast processing time and simplicity.

#### 3) Optimization

Finally, the pipeline was optimized by selecting the best hyper-parameters using a grid-search algorithm. A grid-search algorithm is a brute-force algorithm that searches a user-defined hyper-parameter space to find the most optimal parameters. Here, the YOLO model was used for inference in the event validation set for each study system, and a range of hyper-parameters were defined manually. The hyper-parameters include: YOLO confidence threshold, which is the confidence threshold for a bounding box detection to be considered as a valid detection; minimum duration, which describes the minimum frame number of an event; and 3 separate thresholds for the SORT tracking algorithm, including min hits (minimum frames to define new track), max age (maximum frame gaps to connect two detections) and IOU threshold (intersection over union overlap to associate bounding boxes). These hyper-parameters influence how multiple behavioural detections are grouped together as behavioural events from the SORT tracker. After defining the range of hyper-parameters to explore, we computed the f1-score (see Table 2) for every possible combination of parameters, and then selected the best combination to be used in the final pipeline. In addition, we also determined the best combinations to obtain lowest false negative rates for the event detection case studies, and test whether the framework can be used in hybrid applications. We selected parameters that minimized the false negative rate since that would be the model where the most events were captured for further manual review. We did not optimize for false negative rates in the two frame-wise detection datasets because there is a behaviour prediction every frame, such that a hybrid application will entail reviewing every frame, which is unrealistic.

**Table 2:**
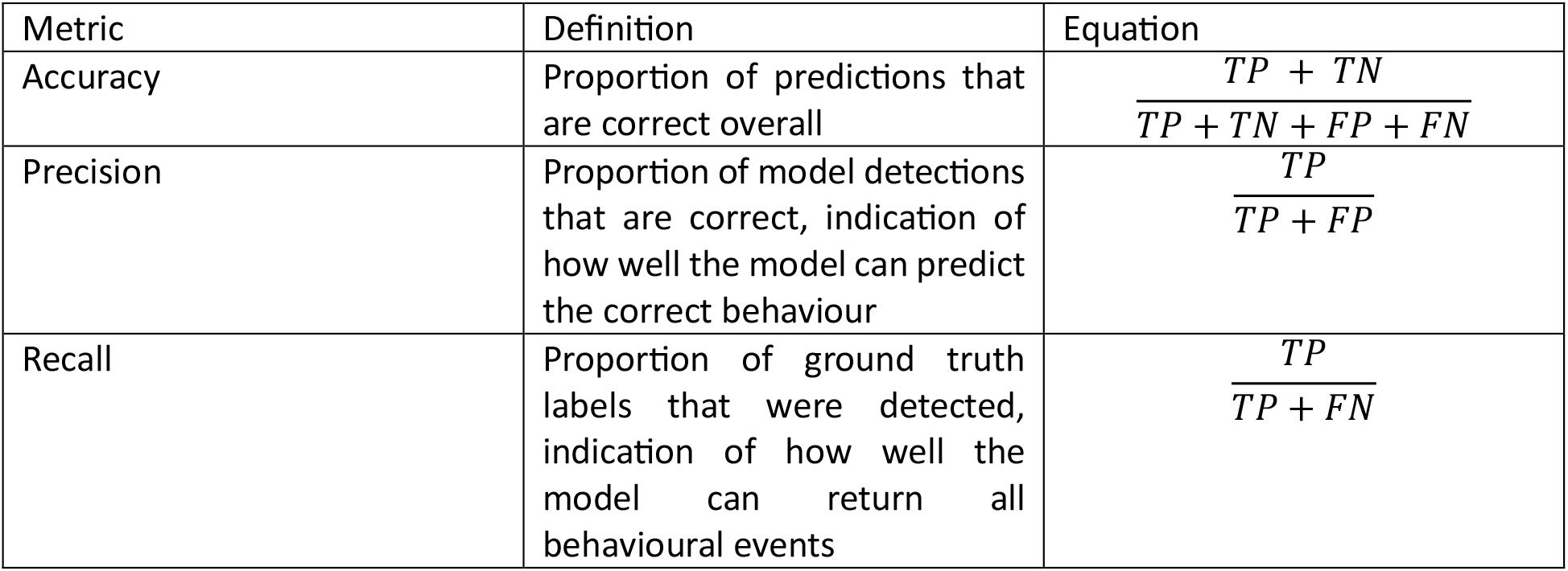
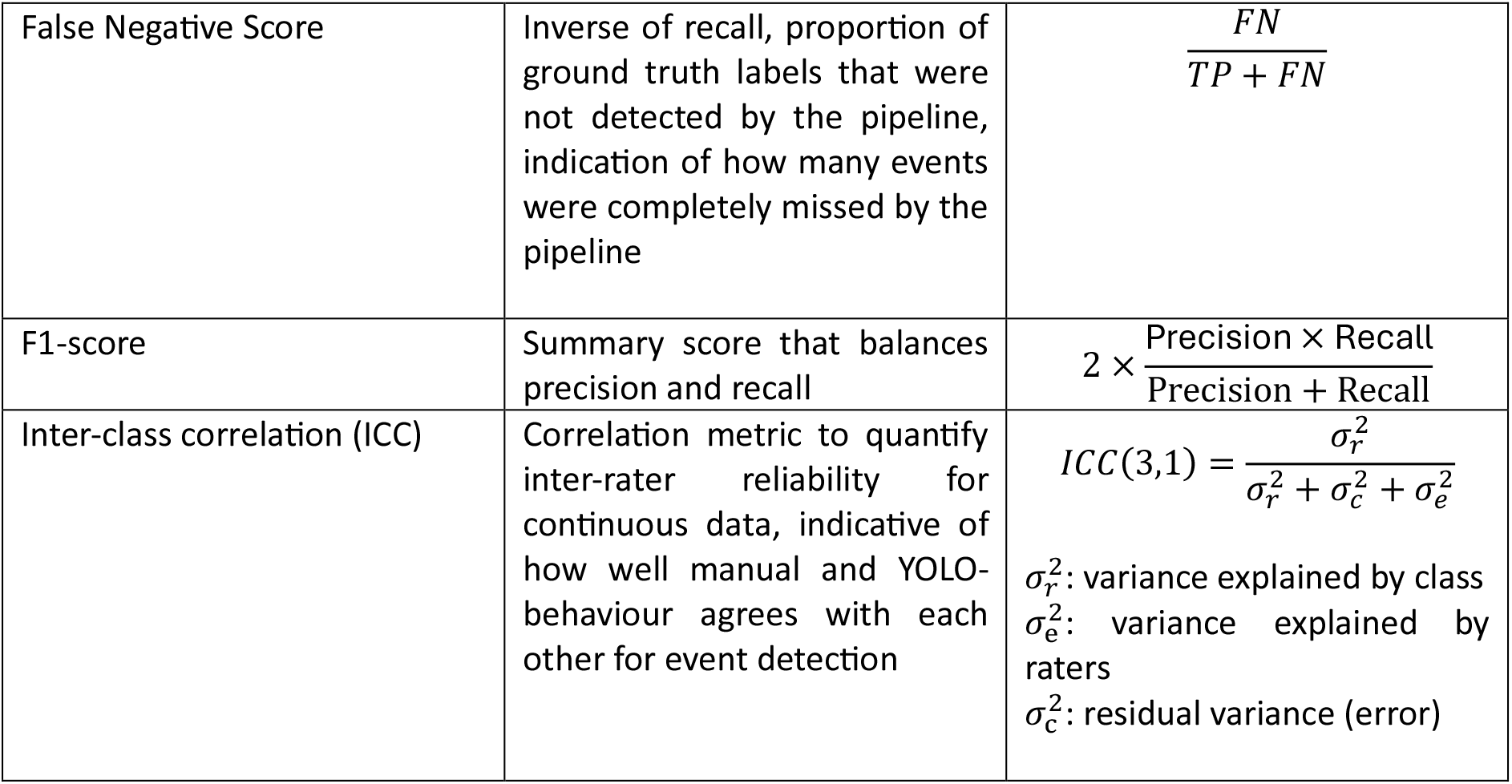
Definitions of evaluation metrics used in the current manuscript. TP: stands for true positives (the model correctly predicts that a given behaviour is present), TN: true negatives (the model correctly predicts that a given behaviour is absent), FP: false positives (the model incorrectly predicts that a given behaviour is present), FN: false negatives (the model incorrectly predicts that a given behaviour is absent), respectively. All metrics ranges from 0 to 1, with higher values representing higher performance

### Evaluation

#### 1) Event Detection

First, we evaluated the reliability of event detection using the event validation dataset for each case study. We used the best parameters obtained from the grid search algorithm for inference, then extracted overall detection accuracy, precision, recall, false negative rate and f1-score (Table 2). Given the disparate characteristics of the datasets, it was necessary to utilise different definitions of “events” for the purposes of evaluation. 1) Sparrow provisioning: each event was defined as a behaviour detection within a 7-second video (see Chan et al., 2024). 2) Jay feeding: each event was defined as 2s windows across the whole video to take into account possible human reaction delay when pressing a button in BORIS, compared to the frame by frame detections of YOLO. Detections were matched as whether a feeding event is present or not within each window. We also report results for a range of time windows to compare how window definition affects evaluation. 3) Human eating: each event was defined as a 1s window due to similar delay in human reaction when coding in BORIS, and detections were matched if an eating event is present within the window. 4) 3D-POP/KABR: Since frame-wise annotations are available, each event is defined as a detection in a single frame

#### 2) Extracting coded behavioural metrics

We used the coded behavioural metrics dataset for each case study to determine whether YOLO-behaviour is reliable for extracting behavioural metrics. For event detection case studies, we extracted feeding rates from jay eating (seconds spent eating per minute per individual) and human eating (food eaten per minute), as well as male and female visit rates (visits per hour) for the sparrow provisioning dataset. For frame-wise detection case studies in 3D-POP and KABR, we extracted the proportion of time spent on each behaviour. We then calculated pearson’s correlation value for each case study, as well as intraclass correlation coefficients (ICC3), which is used to evaluate inter-rater reliability for continuous variables (Gwet, 2014). Since ICC assumes data normality and homogenous variance (Bobak et al., 2018), we log transformed sparrow visit rates and human eating rates, as well as logit transformed proportions from the frame-wise detection datasets and visualizing data distributions to ensure these assumptions were met.

## Results

We applied and evaluated the YOLO-Behaviour framework over all 5 case studies. All YOLO model training evaluation can be found in Supplementary Table 1, and all datasets and annotation used are available here: https://doi.org/10.17617/3.EZNKYV. Qualitative results can also be found in the supplementary video 1.

We found that the YOLO-Behaviour framework is accurate across all study systems and case studies in the event validation set (Table 3), with an f1-score ranging between 0.62 – 0.94 and accuracy ranging between 0.70 - 0.98. Figure 2 shows the confusion matrices for each case study, also highlighting the high accuracy and consistency of most behaviours detected when using YOLO-Behaviour, with little exceptions. Particularly, eating detection for the human dataset is relatively low (0.6, Figure 2C), which is likely due to the inherent difficulty in distinguishing eating from other hand gestures involving the mouth region. In the KABR dataset, the accuracies for locomotion-based behaviours (walking, trotting, running) are more variable (0.27-0.73), as well as browsing and auto-grooming behaviour (0.069, 0.37 respectively, Figure 2E).

**Table 3:**
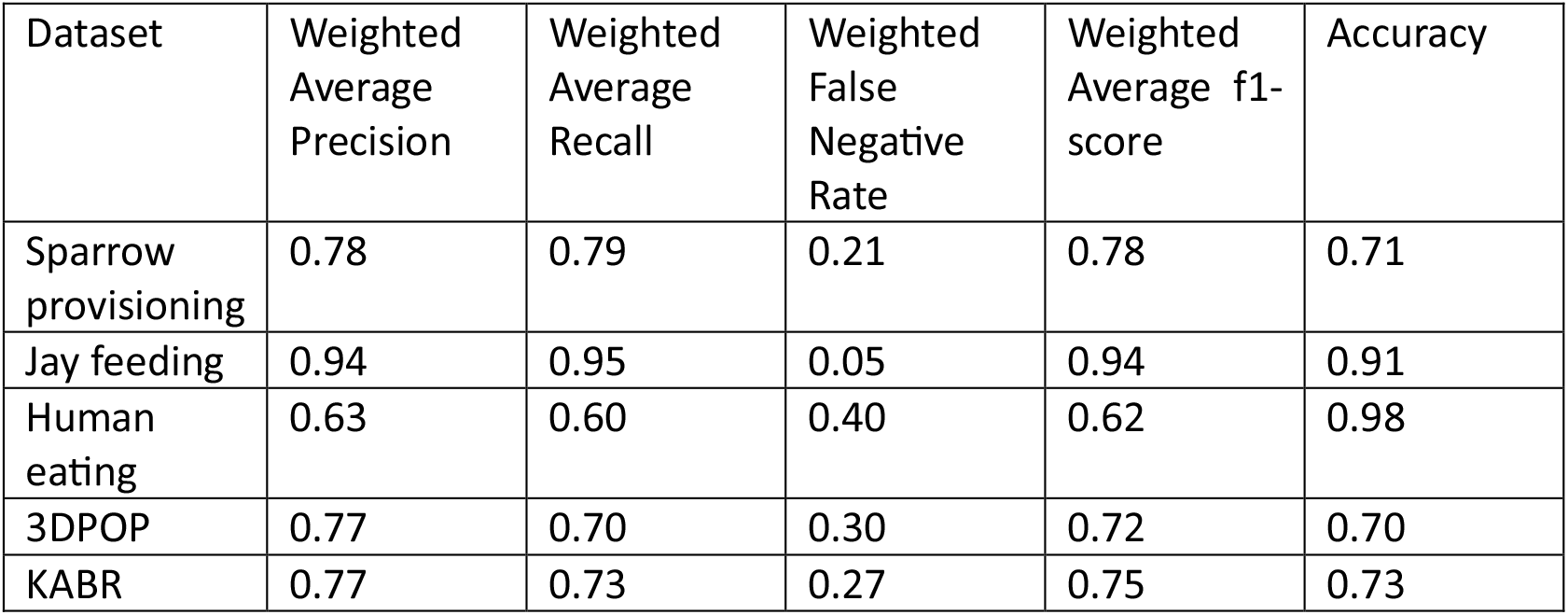
Evaluation metrics on validation set. Presented metrics are the weighted average scores for the behaviour of interest, which excludes “not feeding” for jay and “not eating” for human datasets. For a description of each metric and its definition, we refer to Table 2.

| Dataset | Weighted Average Precision | Weighted Average Recall | Weighted False Negative Rate | Weighted Average f1-score | Accuracy |
| --- | --- | --- | --- | --- | --- |
| Sparrow provisioning | 0.78 | 0.79 | 0.21 | 0.78 | 0.71 |
| Jay feeding | 0.94 | 0.95 | 0.05 | 0.94 | 0.91 |
| Human eating | 0.63 | 0.60 | 0.40 | 0.62 | 0.98 |
| 3DPOP | 0.77 | 0.70 | 0.30 | 0.72 | 0.70 |
| KABR | 0.77 | 0.73 | 0.27 | 0.75 | 0.73 |

**Figure 2:**
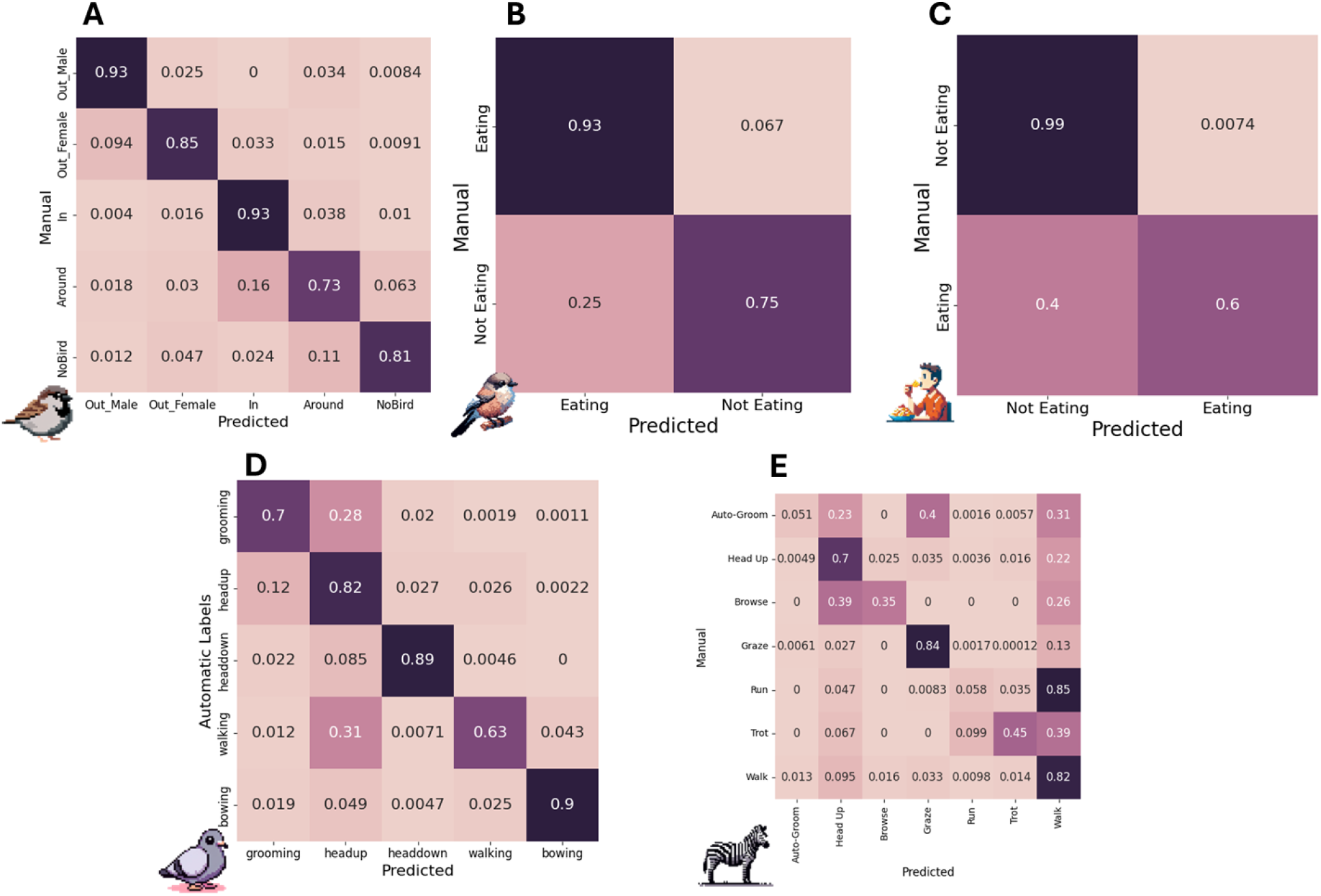
Confusion matrices of per-class classification accuracy across 5 case studies. Accuracy generated from the validation set of each case study, using parameters optimized for f1-score using a grid search algorithm. Values are scaled as the proportion of manual/annotated values. Pixel art of each animal generated using Dall-E 3.

Through the metrics obtained from the coded behavioural metrics dataset, we show that the extracted metrics all significantly correlate with manual annotation (Figure 3), with frame-wise detection case studies having higher correlations and ICC3 values (Table 4; 0.78 – 0.92) compared to event detection (Table 4; 0.49 – 0.70), corresponding to “good” to “excellent” reliability for frame-wise detections and “moderate” to “good” reliability for event-wise detection (Koo and Li, 2016). Figure 3 also shows a general under-detection across all case studies.

**Table 4:**
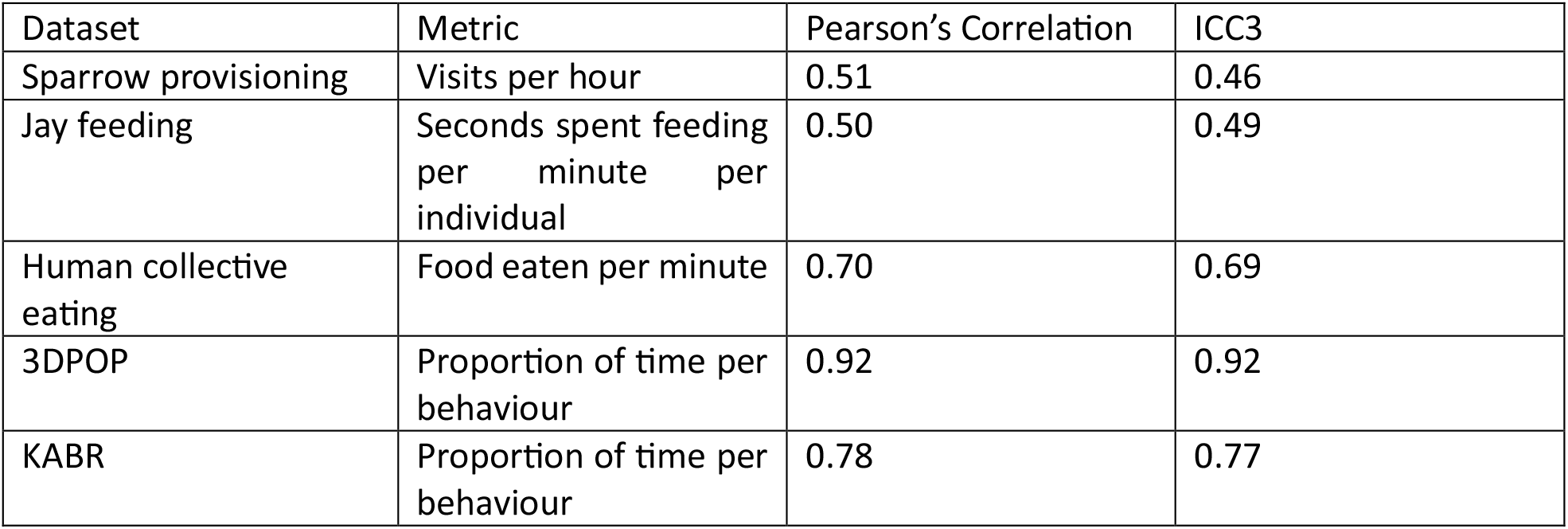
Pearson correlation and Intraclass correlation (ICC3) values for coded behavioural metrics across 5 case studies. Metrics for each case study were extracted from the coded behavioural metrics dataset. We refer to Table 1 for description of the datasets and Table 2 for the definition of ICC3. All Pearson’s correlation values were significant (p< 0.05)

| Dataset | Metric | Pearson's Correlation | ICC3 |
| --- | --- | --- | --- |
| Sparrow provisioning | Visits per hour | 0.51 | 0.46 |
| Jay feeding | Seconds spent feeding per minute per individual | 0.50 | 0.49 |
| Human collective eating | Food eaten per minute | 0.70 | 0.69 |
| 3DPOP | Proportion of time per behaviour | 0.92 | 0.92 |
| KABR | Proportion of time per behaviour | 0.78 | 0.77 |

**Figure 3:**
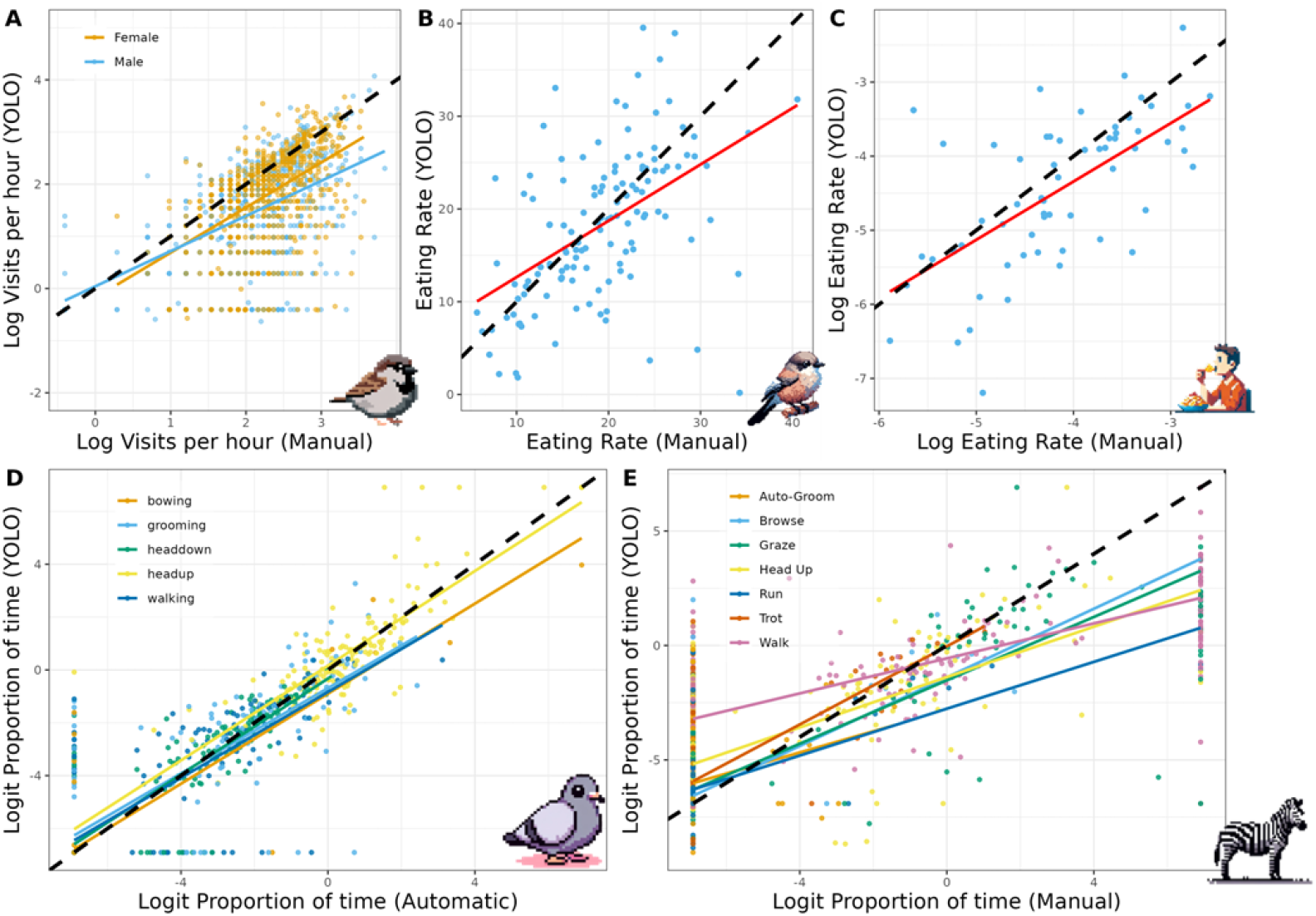
Correlations of the coded behavioural metrics across 5 case studies. Metrics were extracted from the coded behavioural metrics dataset of each case study. The y-axis showing automatic rates from YOLO and the x-axis showing rates from manual or labels from corresponding datasets. Black line in each plot represents the 1:1 correlation line. Correlation of visit rates (visits per hour) in the sparrow provisioning dataset, separated by male and female visit rates. B) Correlation of feeding rate (seconds spent feeding per minute per individual) in the Siberian jay dataset. C) Correlation of eating rate (food eaten per minute) of the human eating dataset. D) Correlation of the proportion of time spent for five separate behaviours in the pigeon 3D-POP dataset. E) Correlation of the proportion of time spent for seven separate behaviours in the Giraffe/Zebra KABR dataset. Pixel art of each animal generated using Dall-E 3. A)

Finally, we tested whether the YOLO-Behaviour framework can be used in a hybrid approach by optimizing for low false negative rates instead of f1-score. We found that the method can obtain low false negative rates between 0.05 – 0.14 (Table 5), showing that only around 5% to 15% of the overall events will be missed. Particularly in the human eating case study, the precision is 0.26, which shows that the model is full of false positives that can be manually corrected in the hybrid framework.

**Table 5:** Evaluation metrics from the validation set, optimized by recall for event detection case studies. Metrics were optimized for recall instead of f1-score during the grid search algorithm, to test whether the framework can be used as a hybrid method.

| Dataset | Average Precision | Average Recall | Average False Negative Rate | Average f1-score | Accuracy |
| --- | --- | --- | --- | --- | --- |
| Sparrow provisioning | 0.78 | 0.87 | 0.13 | 0.82 | 0.71 |
| Jay feeding | 0.94 | 0.95 | 0.05 | 0.95 | 0.91 |
| Human collective eating | 0.26 | 0.86 | 0.14 | 0.40 | 0.94 |

## Discussion

In the current study, we presented the YOLO-Behaviour framework, a simple to implement and flexible framework for automated coding for animal behaviour from videos. We illustrated the robustness of the framework with the high detection accuracy across a large range of study systems and video types. The framework is easy to train and implement, with the full documentation and user guidelines for applying to a new system available here: https://alexhang212.github.io/YOLO_Behaviour_Repo/.

Using the YOLO-behaviour framework, we contributed to efforts towards quantifying behavioural metrics across 5 diverse study systems. For example, in the Lundy house sparrow system, we obtained parental provisioning rates for more than double the sample size compared to manual annotated data (Chan et al., 2024), allowing for stronger basis to discover the drivers and consequences of parental care behaviour, including indirect genetic effects (Schroeder et al., 2019) or its fitness outcomes (Schroeder et al., 2013). In Siberian jays, the method can be applied across the huge backlog of videos to gain insight into the co-feeding behaviour of this family living species where groups also can include unrelated non-breeders (Griesser, 2003; Griesser et al., 2015), to reveal the mechanisms facilitating the evolution of breeding systems and cooperation (Drobniak et al., 2015). In the human eating dataset, while the current dataset was obtained from a specific experiment, future data collection can benefit from the framework to reduce annotation time. Finally, while the 3D-POP and KABR datasets were used from computer vision datasets to demonstrate the ability for the framework to do frame-wise detections, the framework shows promise for larger scale deployment to quantify activity budgets of animals from drones (Koger et al., 2023; Schad and Fischer, 2023) or from citizen science data.

### Evaluation

The framework was first evaluated via the validation set for each case study to determine its accuracy (in terms of precision) and its capacity to return a majority of the manually coded events (in terms of recall). Overall, the framework performed well for all study systems with little exceptions. Firstly, the precision-recall for the human eating dataset was relatively low (around 0.6), which can be caused by hand gestures near the participant’s mouth region, which is visually similar to the eating behaviour. To address this issue, additional training data might need to be added, or a hybrid human-in-the-loop method can be considered (see below). For the KABR dataset, we identified a few behaviours that the framework could not detect accurately. The first behaviour concerns locomotion, including run, trot and walk, which were difficult to distinguish without temporal context, since YOLO is a frame-wise method. For deployment of the model to detect these locomotion behaviours, a possible solution can be to add a speed threshold to separate the 3 behaviours, which cannot be tested in the current manuscript since the animal subjects in the KABR dataset were always centered in frame (Kholiavchenko et al., 2024). We note that auto-grooming and browsing behaviour was also not well detected using the YOLO-Behaviour framework (Figure 2E), which was probably caused by the large variation in postures/ visual appearances of both behaviours in zebras and giraffes, making the model unable to generalize beyond the training data.

While precision recall values were high for the Siberian jay feeding dataset, we note that we used a 2s time window to match behaviours, as we observed a large mismatch between human annotation and automated detection. We also report results for different time window intervals (Supplementary Table 3), and we show that all evaluation metrics improve up until the 2s time window, which we assume is the appropriate window to match up automated detection and manual annotation. This mismatch can be caused by human reaction delay, especially when videos were coded in real time, but can also be due to incorrect detections by the model. However, upon visually inspecting qualitative results (Supplementary video 1), it seems more likely that there is a mismatch in frame-by-frame detections between automated and human manual annotations, which is important to consider when evaluating machine learning models in future studies.

Next, we used the coded behavioural metrics dataset to test whether the YOLO-Behaviour framework can be used to extract behavioural metrics. Overall, we found high correlations and ICC values for both pigeon and zebra/giraffe frame-wise detections, but lower values for the other event-wise detection case studies. For human eating detection, this was expected due to the low evaluation metrics from the event-wise validation, and the low correlation can be the effect of misidentification of the eating behaviour itself. However, for the jay eating and sparrow provisioning datasets, we found low correlation albeit high precision-recall metrics from the event validation. For the jay dataset, this can be attributed to a mismatch between manual annotation of feeding behaviour in BORIS and the automatic method, and for the sparrow dataset, this can be caused by the many observers who annotated the dataset over the years that can result in inconsistent visit rates. We also note that there is a general under-detection of coded behavioural metrics extracted by YOLO-behaviour across datasets, meaning false negative rates (missed detections) were a stronger contributor to the low accuracy, which can potentially be improved with additional training data. Finally, we acknowledge that the correlation and ICC values might not be directly indicative of how well the YOLO-Behaviour framework can predict coded behavioural metrics, and additional evaluation like hypothesis testing (see Chan et al., 2024) could be useful to further validate the method.

### Hybrid applications

In cases where automated detection accuracy might be low and insufficient for a certain study system, we also tested whether the YOLO-Behaviour framework can be used as a pre-processing step in a hybrid approach. Instead of optimizing for f1-score, we optimized for low false negative rates using grid search and found very low false negatives across all event detection datasets. For example, in the human eating dataset, we retrieved low false negative rates (0.14) and high recall (0.86) by trading off low precision (0.26). Hence, by first using YOLO-behaviour to extract events then manually confirm whether the detections were correct, we can potentially reach up to 0.86 precision in human eating detection, compared to the 0.63 precision when using the framework in a fully automated manner. While some manual annotation is still required, this hybrid approach would further reduce annotation time with the assurance that extracted behavioural events are accurate. In the provided code-base, we also provide example code to run the pipeline with a human–in-the-loop approach.

### Limitations

The YOLO model used in the current framework only takes a single frame as input, which might not be able to reliably detect behaviours that have a temporal aspect, like the locomotion behaviours in KABR. However, we do note that the current framework was able to detect walking and bowing behaviours in the pigeon dataset reliably, likely due to other visual cues (e.g., leg up, puffed up neck). For behaviours that have an important temporal component, other methods that take video input (Rodríguez-Moreno et al., 2019), or first do posture estimation (Mathis and Mathis, 2020) might be considered. Still, we note that compared to the 86.7% per-instance accuracy reported in the original KABR publication (Kholiavchenko et al., 2024) using a temporal based X3D method, the current YOLO-Behaviour framework still managed to recover similar accuracies (72%, Table 3)., Whether this difference in accuracy is important for quantifying behaviours will depend on the specific use case.

Finally, similar to any commonly used deep learning model is the requirement for training data. While the case studies in the current paper only used minimal training data (∽1000-1500 images), the framework might not work too well if the behaviour of interest is very rare (only a few instances available). In these instances other methods like few-shot object detection models (Köhler et al., 2023) might be considered a better fit. However, compared to methods that first do posture estimation and then behavioural recognition, our current framework saves a large amount of required training data by only requiring behavioural annotations within a frame, compared to posture annotations, as well as time-series annotations on behavioural sequences (e.g., Hu et al., 2023; Wittek et al., 2022). We also highlight that in practice, any object detector model can also be used in the same way to quantify behaviour, but we chose YOLO here because of the ease of use and robustness.

### Conclusion

In conclusion, we presented the YOLO-Behaviour Framework, a simple, flexible and robust method for automating video annotation of behaviours. We demonstrated the efficiency of the pipeline in 5 distinct case studies, and highlighted that the framework works well across a wide range of behaviours and videos. With the increased use of deep learning and machine learning for measuring behaviours in animals, we hope the framework can be another step towards lowering the barrier to train and deploy these methods, and replacing time-consuming manual annotation in the field of behaviour research.

## Acknowledgements

This work is funded by the Deutsche Forschungsgemeinschaft (DFG, German Research Foundation) under Germany’s Excellence Strategy – EXC 2117 – 422037984. We additionally thank Francesca Frisoni for additional manual annotation, and all students/ field assistants who collected and manually annotated the datasets over the years. I also thank Stephen Tyndel for sharing my enthusiasm on the wonders of YOLO and commenting on an initial draft of the manuscript. MG was supported by a Heisenberg Grant no. GR 4650/2-1 by the German Research Foundation DFG.

## Author Contributions

AHHC, PP, HS conceived the ideas and designed methodology; HS,JK,JSr,BR,JSc,WDP,SN,TB,MG,AM,SL collected the data; AHHC analysed the data; AHHC led the writing of the manuscript. FK supervised the project. All authors contributed critically to the drafts and gave final approval for publication.

## References

Bewley, A., Ge, Z., Ott, L., Ramos, F., Upcroft, B., 2016. Simple Online and Realtime Tracking, in: 2016 IEEE International Conference on Image Processing (ICIP). pp. 3464–3468. 10.1109/ICIP.2016.7533003

Bobak, C.A., Barr, P.J., O’Malley, A.J., 2018. Estimation of an inter-rater intra-class correlation coefficient that overcomes common assumption violations in the assessment of health measurement scales. BMC Med. Res. Methodol. 18, 93. 10.1186/s12874-018-0550-6

Brookes, O., Mirmehdi, M., Stephens, C., Angedakin, S., Corogenes, K., Dowd, D., Dieguez, P., Hicks, T.C., Jones, S., Lee, K., Leinert, V., Lapuente, J., McCarthy, M.S., Meier, A., Murai, M., Normand, E., Vergnes, V., Wessling, E.G., Wittig, R.M., Langergraber, K., Maldonado, N., Yang, X., Zuberbühler, K., Boesch, C., Arandjelovic, M., Kühl, H., Burghardt, T., 2024. PanAf20K: A Large Video Dataset for Wild Ape Detection and Behaviour Recognition. Int. J. Comput. Vis. 10.1007/s11263-024-02003-z

Chan, A.H.H., Liu, J., Burke, T., Pearse, W.D., Schroeder, J., 2024. Comparison of manual, machine learning, and hybrid methods for video annotation to extract parental care data. J. Avian Biol. 2024, e03167. 10.1111/jav.03167

Chen, J., Hu, M., Coker, D.J., Berumen, M.L., Costelloe, B., Beery, S., Rohrbach, A., Elhoseiny, M., 2023. MammalNet: A Large-Scale Video Benchmark for Mammal Recognition and Behavior Understanding. Presented at the Proceedings of the IEEE/CVF Conference on Computer Vision and Pattern Recognition, pp. 13052–13061.

Chimento, M., Chan, A.H.H., Aplin, L.M., Kano, F., 2024. Peering into the world of wild passerines with 3D-SOCS: synchronized video capture and posture estimation. bioRxiv 2024.06.30.601375. 10.1101/2024.06.30.601375

Christin, S., Hervet, É., Lecomte, N., 2019. Applications for deep learning in ecology. Methods Ecol. Evol. 10, 1632–1644.

Couzin, I.D., Heins, C., 2022. Emerging technologies for behavioral research in changing environments. Trends Ecol. Evol. 10.1016/j.tree.2022.11.008

Delacoux, M., Kano, F., 2024. Fine-scale tracking reveals visual field use for predator detection and escape in collective foraging of pigeon flocks. 10.1101/2024.02.05.578919

Dell, A.I., Bender, J.A., Branson, K., Couzin, I.D., de Polavieja, G.G., Noldus, L.P., Pérez-Escudero, A., Perona, P., Straw, A.D., Wikelski, M., 2014. Automated image-based tracking and its application in ecology. Trends Ecol. Evol. 29, 417–428.

Drobniak, S.M., Wagner, G., Mourocq, E., Griesser, M., 2015. Family living: an overlooked but pivotal social system to understand the evolution of cooperative breeding. Behav. Ecol. 26, 805–811. 10.1093/beheco/arv015

Friard, O., Gamba, M., 2016. BORIS: a free, versatile open-source event-logging software for video/audio coding and live observations. Methods Ecol. Evol. 7, 1325–1330. 10.1111/2041-210X.12584

Goss, K., Bueno-Junior, L.S., Stangis, K., Ardoin, T., Carmon, H., Zhou, J., Satapathy, R., Baker, I., Jones-Tinsley, C.E., Lim, M.M., Watson, B.O., Sueur, C., Ferrario, C.R., Murphy, G.G., Ye, B., Hu, Y., 2024. Quantifying social roles in multi-animal videos using subject-aware deep-learning. BioRxiv Prepr. Serv. Biol. 2024.07.07.602350. 10.1101/2024.07.07.602350

Graving, J.M., Chae, D., Naik, H., Li, L., Koger, B., Costelloe, B.R., Couzin, I.D., 2019. DeepPoseKit, a software toolkit for fast and robust animal pose estimation using deep learning. Elife 8, e47994.

Graving, J.M., Couzin, I.D., 2020. VAE-SNE: a deep generative model for simultaneous dimensionality reduction and clustering. 10.1101/2020.07.17.207993

Griesser, M., 2003. Nepotistic vigilance behavior in Siberian jay parents. Behav. Ecol. 14, 246–250. 10.1093/beheco/14.2.246

Griesser, M., Halvarsson, P., Drobniak, S.M., Vilà, C., 2015. Fine-scale kin recognition in the absence of social familiarity in the Siberian jay, a monogamous bird species. Mol. Ecol. 24, 5726–5738. 10.1111/mec.13420

Gwet, K.L., 2014. Handbook of Inter-Rater Reliability, 4th Edition: The Definitive Guide to Measuring The Extent of Agreement Among Raters. Advanced Analytics, LLC.

Hsu, A.I., Yttri, E.A., 2021. B-SOiD, an open-source unsupervised algorithm for identification and fast prediction of behaviors. Nat. Commun. 12, 5188. 10.1038/s41467-021-25420-x

Hu, Y., Ferrario, C.R., Maitland, A.D., Ionides, R.B., Ghimire, A., Watson, B., Iwasaki, K., White, H., Xi, Y., Zhou, J., Ye, B., 2023. LabGym: Quantification of user-defined animal behaviors using learning-based holistic assessment. Cell Rep. Methods 3, 100415. 10.1016/j.crmeth.2023.100415

Jocher, G., Chaurasia, A., Qiu, J., 2023. YOLO by Ultralytics.

Joska, D., Clark, L., Muramatsu, N., Jericevich, R., Nicolls, F., Mathis, A., Mathis, M.W., Patel, A., 2021. AcinoSet: A 3D Pose Estimation Dataset and Baseline Models for Cheetahs in the Wild, in: 2021 IEEE International Conference on Robotics and Automation (ICRA). pp. 13901–13908. 10.1109/ICRA48506.2021.9561338

Kholiavchenko, M., Kline, J., Ramirez, M., Stevens, S., Sheets, A., Babu, R., Banerji, N., Campolongo, E., Thompson, M., Van Tiel, N., Miliko, J., Bessa, E., Duporge, I., Berger-Wolf, T., Rubenstein, D., Stewart, C., 2024. KABR: In-Situ Dataset for Kenyan Animal Behavior Recognition from Drone Videos, in: 2024 IEEE/CVF Winter Conference on Applications of Computer Vision Workshops (WACVW). Presented at the 2024 IEEE/CVF Winter Conference on Applications of Computer Vision Workshops (WACVW), IEEE, Waikoloa, HI, USA, pp. 31–40. 10.1109/WACVW60836.2024.00011

Koger, B., Deshpande, A., Kerby, J.T., Graving, J.M., Costelloe, B.R., Couzin, I.D., 2023. Quantifying the movement, behaviour and environmental context of group-living animals using drones and computer vision. J. Anim. Ecol. 92, 1357–1371. 10.1111/1365-2656.13904

Köhler, M., Eisenbach, M., Gross, H.-M., 2023. Few-Shot Object Detection: A Comprehensive Survey. IEEE Trans. Neural Netw. Learn. Syst. 1–21. 10.1109/TNNLS.2023.3265051

Koo, T.K., Li, M.Y., 2016. A Guideline of Selecting and Reporting Intraclass Correlation Coefficients for Reliability Research. J. Chiropr. Med. 15, 155–163. 10.1016/j.jcm.2016.02.012

Lei, Y., Dong, P., Guan, Y., Xiang, Y., Xie, M., Mu, J., Wang, Y., Ni, Q., 2022. Postural behavior recognition of captive nocturnal animals based on deep learning: a case study of Bengal slow loris. Sci. Rep. 12, 7738. 10.1038/s41598-022-11842-0

Lin, T.-Y., Maire, M., Belongie, S., Hays, J., Perona, P., Ramanan, D., Dollár, P., Zitnick, C.L., 2014. Microsoft coco: Common objects in context, in: European Conference on Computer Vision. Springer, pp. 740–755.

Mathis, A., Mamidanna, P., Cury, K.M., Abe, T., Murthy, V.N., Mathis, M.W., Bethge, M., 2018. DeepLabCut: markerless pose estimation of user-defined body parts with deep learning. Nat. Neurosci.

Mathis, M.W., Mathis, A., 2020. Deep learning tools for the measurement of animal behavior in neuroscience. Curr. Opin. Neurobiol. 60, 1–11.

Naik, H., Chan, A.H.H., Yang, J., Delacoux, M., Couzin, I.D., Kano, F., Nagy, M., 2023. 3D-POP - An Automated Annotation Approach to Facilitate Markerless 2D-3D Tracking of Freely Moving Birds With Marker-Based Motion Capture. Presented at the Proceedings of the IEEE/CVF Conference on Computer Vision and Pattern Recognition, pp. 21274–21284.

Nakagawa, S., Gillespie, D.O.S., Hatchwell, B.J., Burke, T., 2007. Predictable males and unpredictable females: sex difference in repeatability of parental care in a wild bird population. J. Evol. Biol. 20, 1674–1681.

Ng, X.L., Ong, K.E., Zheng, Q., Ni, Y., Yeo, S.Y., Liu, J., 2022. Animal Kingdom: A Large and Diverse Dataset for Animal Behavior Understanding. Presented at the Proceedings of the IEEE/CVF Conference on Computer Vision and Pattern Recognition, pp. 19023–19034.

Pereira, T.D., Tabris, N., Matsliah, A., Turner, D.M., Li, J., Ravindranath, S., Papadoyannis, E.S., Normand, E., Deutsch, D.S., Wang, Z.Y., McKenzie-Smith, G.C., Mitelut, C.C., Castro, M.D., D’Uva, J., Kislin, M., Sanes, D.H., Kocher, S.D., Wang, S.S.-H., Falkner, A.L., Shaevitz, J.W., Murthy, M., 2022. SLEAP: A deep learning system for multi-animal pose tracking. Nat. Methods 19, 486–495. 10.1038/s41592-022-01426-1

Rodríguez-Moreno, I., Martínez-Otzeta, J.M., Sierra, B., Rodriguez, I., Jauregi, E., 2019. Video Activity Recognition: State-of-the-Art. Sensors 19, 3160. 10.3390/s19143160

Schad, L., Fischer, J., 2023. Opportunities and risks in the use of drones for studying animal behaviour. Methods Ecol. Evol. 14, 1864–1872. 10.1111/2041-210X.13922

Schroeder, J., Cleasby, I., Dugdale, H.L., Nakagawa, S., Burke, T., 2013. Social and genetic benefits of parental investment suggest sex differences in selection pressures. J. Avian Biol. 44, 133–140.

Schroeder, J., Dugdale, H., Nakagawa, S., Sparks, A., Burke, T., 2019. Social genetic effects (IGE) and genetic intra-and intersexual genetic correlation contribute to the total heritable variance in parental care.

Schroeder, J., Hsu, Y.-H., Winney, I., Simons, M., Nakagawa, S., Burke, T., 2020. Data from: Predictably philandering females prompt poor paternal provisioning. 10.5061/DRYAD.0T313

Tuia, D., Kellenberger, B., Beery, S., Costelloe, B.R., Zuffi, S., Risse, B., Mathis, A., Mathis, M.W., van Langevelde, F., Burghardt, T., 2022. Perspectives in machine learning for wildlife conservation. Nat. Commun. 13, 1–15.

Tuyttens, F.A.M., de Graaf, S., Heerkens, J.L., Jacobs, L., Nalon, E., Ott, S., Stadig, L., Van Laer, E., Ampe, B., 2014. Observer bias in animal behaviour research: can we believe what we score, if we score what we believe? Anim. Behav. 90, 273–280.

Waldmann, U., Chan, A.H.H., Naik, H., Nagy, M., Couzin, I.D., Deussen, O., Goldluecke, B., Kano, F., 2024. 3D-MuPPET: 3D Multi-Pigeon Pose Estimation and Tracking. Int. J. Comput. Vis. 10.1007/s11263-024-02074-y

Weinstein, B.G., 2018. A computer vision for animal ecology. J. Anim. Ecol. 87, 533–545.

Wittek, N., Wittek, K., Keibel, C., Güntürkün, O., 2022. Supervised machine learning aided behavior classification in pigeons. Behav. Res. Methods. 10.3758/s13428-022-01881-w

Yang, A., Huang, H., Yang, X., Li, S., Chen, C., Gan, H., Xue, Y., 2019. Automated video analysis of sow nursing behavior based on fully convolutional network and oriented optical flow. Comput. Electron. Agric. 167, 105048.

