## SupplementaryMaterials for "YOLO-Behaviour: A simple, flexible framework to automatically quantify animal behaviours from videos"

**Supplementary Materials**

Supplementary Methods

Datasets

1) House sparrow provisioning

The house sparrow (*Passer domesticus*) provisioning dataset is collected from a long-term study system on Lundy Island, UK (51°10'N, 4°40'W), where life history, genetics and breeding were monitored over the last 20+ years (Schroeder et al., 2020).As part of the yearly monitoring protocol, 90-minute long provisioning videos (1280x720, 25fps) were recorded on the 7th and 14th day of egg hatching for each breeding pair over the breeding season to quantify parental investment rates (Chan et al., 2024; Nakagawa et al., 2007). From this, we manually annotated 4 behaviours: Male out, Female out, In and Around. Since house sparrows expresses sexual dimorphism, with the throat badge being the most prevalent feature distinguishing male and female sparrows, we annotated the sex only when birds are flying out of the nest box, since the black badge is not visible when birds are entering the nest box. To calculate final male and female visit rates, we tallied the number of Male out and Female out events and divided by the total length to get visits per hour.

2) Siberian jay feeding

The Siberian jay (*Perisoreus infaustus*) feeding dataset were collected from a long term study population northwest of Arvidsjaur, northern Sweden (65°40' N, 19°0' E) in autumn 2021. Siberian Jays are a species that form territorial and cooperative groups that are made of kin and non-kin individuals. Each autumn, 15-30 minute long behavioural videos were repeatedly taken of a wooden perch placed in each territory, with pieces of fat in the middle, to attract arriving birds. From this, dominance relationships, feeding rate and submissive behaviour were manually coded using BORIS. For the event validation dataset, we additionally annotated 5 videos in finer temporal scales, by marking each time a bird was pecking the fat. In the YOLO-behaviour framework, we labelled frames where the birds were pecking at the fat, by drawing a bounding box that includes both the beak and fat. We did not detect submissive behaviour, since the behaviour was too short and similar to flight. We also did not detect agonistic behaviours (e.g displacement), since that would require automated identification of individuals, which is outside the scope of this study.

3) Human collective eating dataset

The human (*Homo sapiens*) collective eating dataset is part of the collective appetite project at the Centre for the Advanced Study of Collective Behaviour that aims to determine how eating together influences human relationships, behaviour, and experiences.. For this analysis data captured by 3 cameras, one aimed at each participant, and later cropped to only include the focal individual was used. Behaviours for each video were manually coded using BORIS, which includes eating, dipping and taking nacho chips. For YOLO-Behaviour, only eating was selected, since taking and dipping behaviour is often mis-detected, due to other participants reaching for the nachos and dipping bowls. Due to privacy and ethical reasons, no image or video data from the experiment can be shared.

4) 3D-Posture of Pigeons (3D-POP)

The homing pigeon (*Colomba livia*) dataset is part of the 3D Postures of pigeons (3D-POP) dataset. 3D-POP is a large-scale, multi-view, multi-individual 2D to 3D dataset for object detection, posture estimation and tracking (Naik et al., 2023), which includes sequences of 1,2,5 and 10 pigeons. The dataset is constructed with a semi-automatic pipeline using a marker-based motion capture system. However, the dataset does not include behavioural annotations, hence we used the raw 3D motion capture data to automatically classify behaviours based on multiple thresholds. We refer to the supplementary materials of Delacoux and Kano (2024) for detailed description and validation of the automatic behavioural classification. Using this automated behavioural coding, we then combined 5 distinct behaviours (walking, head-up, head-down, grooming and bowing) with the ground truth bounding box extracted from 3D-POP as training data for YOLO-Behaviour.

5) In-situ dataset for Kenyan animal behaviour recognition (KABR)

The zebra/ giraffe dataset is part of the in-situ dataset for Kenyan animal behaviour recognition (KABR), which is a comprehensive behavioural dataset of drone videos filming Giraffes (*Giraffa camelopardalis*), plain zebras (*Equus quagga*) and Grevy's zebras (*Equus grevyi*) from the Mpala research centre, Kenya. The KABR dataset provides "mini-scenes" with the resolution of 400x300px, which includes an animal centered at the middle of the scene, and frame by frame behavioural annotations. However, the dataset does not include bounding box annotations. To obtain bounding box annotations to train the YOLO-behaviour framework, we used a pre-trained YOLOv8l model, which includes both "giraffe" and "zebra" as available classes based on the COCO dataset (Lin et al., 2014). We then assigned each bounding box to the behaviour of each given frame, and take the bounding box closest to the mid-point if multiple giraffe or zebras were detected. After this procedure, we then have "mini-scenes" with bounding box and behavioural annotations for 7 distinct behaviours (walk, graze, browse, head-up, groom, trot, run). Note that we removed the class "occluded" from the KABR dataset to avoid mis-leading or confusing the YOLO-behaviour model.

**Supplementary Results**

**Supplementary Table 1: YOLO evaluation metrics from the training data test set**

| Dataset | Precision | Recall | mAP50 |
|---|---|---|---|
| Sparrow | 0.97 | 0.92 | 0.97 |
| Jay | 0.80 | 0.80 | 0.87 |
| Human | 1 | 1 | 0.995 |
| Pigeon (3D-POP) | 0.63 | 0.79 | 0.77 |
| Zebra/Giraffe (KABR) | 0.88 | 0.91 | 0.94 |

**Supplementary Table 2:** Final hyper parameters used for evaluation after optimizing for the highest f1-score from the grid search algorithm

| Dataset | Behaviour | Max age | Min hit | Iou threshold | Min duration | YOLO threshold |
|---|---|---|---|---|---|---|
| Sparrow Provisioning | Out_Male | 1 | 2 | 0.1 | 7 | 0.5 |
| | Out_Female | 1 | 3 | 0.1 | 4 | 0.5 |
| | In | 1 | 0 | 0.2 | 5 | 0.4 |
| | Around | 6 | 0 | 0.4 | 9 | 0.3 |
| Jay Eating | Eat | 21 | 1 | 0.2 | 1 | 0.1 |
| Human Eating | Eat | 6 | 0 | 0 | 4 | 0.6 |
| Pigeon (3D-POP) | Head-up | 1 | 3 | 0.5 | 10 | 0.3 |
| | Head-down | 6 | 1 | 0.5 | 10 | 0.5 |

| | | | | | | |
|---|---|---|---|---|---|---|
| | Walking | 6 | 3 | 0.5 | 20 | 0.3 |
| | Bowing | 11 | 5 | 0.3 | 20 | 0.3 |
| | Grooming | 21 | 3 | 0.7 | 10 | 0.3 |
| Zebra, Giraffe (KABR) | Auto-Groom | 1 | 9 | 0.1 | 10 | 0.8 |
| | Head up | 6 | 1 | 0.1 | 20 | 0 |
| | Browse | 11 | 9 | 0.1 | 20 | 0 |
| | Graze | 21 | 1 | 0.1 | 15 | 0 |
| | Run | 1 | 5 | 0.1 | 20 | 0.7 |
| | Trot | 1 | 9 | 0.1 | 10 | 0.7 |
| | Walk | 16 | 1 | 0.1 | 15 | 0 |

**Supplementary Table 3:** Final hyper parameters used for evaluation after optimizing for the highest recall score from the grid search algorithm, for event detection case studies

| Dataset | Behaviour | Max age | Min hit | Iou threshold | Min duration | YOLO threshold |
|---|---|---|---|---|---|---|
| Sparrow Provisioning | Out_Male | 1 | 2 | 0.1 | 7 | 0.5 |
| | Out_Female | 1 | 3 | 0.1 | 4 | 0.5 |
| | In | 1 | 0 | 0.2 | 5 | 0.4 |
| | Around | 6 | 0 | 0.4 | 9 | 0.3 |
| Jay Eating | Eat | 21 | 1 | 0.1 | 1 | 0.1 |
| Human Eating | Eat | 21 | 0 | 0 | 0 | 0 |

**Supplementary Table 4:** Evaluation metrics for different time window choice for the Siberian jay dataset, highlighting increasing accuracy as time window is increased.

| Time interval (frames) | Average Precision | Average Recall | Average f1-score | Cohen's Kappa | Accuracy |
|---|---|---|---|---|---|
| 1 frame, 1/25 seconds | 0.03 | 0.29 | 0.06 | 0.01 | 0.74 |
| 13 frames, ~0.52 seconds | 0.41 | 0.62 | 0.50 | 0.13 | 0.56 |
| 25 frames, 1 second | 0.70 | 0.82 | 0.75 | 0.32 | 0.68 |
| 50 frames, 2 seconds | 0.94 | 0.95 | 0.94 | 0.73 | 0.91 |
| 75 frames, 3 seconds | 0.98 | 0.98 | 0.98 | 0.87 | 0.96 |

**References**

Chan, A.H.H., Liu, J., Burke, T., Pearse, W.D., Schroeder, J., 2024. Comparison of manual, machine learning, and hybrid methods for video annotation to extract parental care data. J. Avian Biol. 2024, e03167. https://doi.org/10.1111/jav.03167

Delacoux, M., Kano, F., 2024. Fine-scale tracking reveals visual field use for predator detection and escape in collective foraging of pigeon flocks. https://doi.org/10.1101/2024.02.05.578919

Lin, T.-Y., Maire, M., Belongie, S., Hays, J., Perona, P., Ramanan, D., Dollár, P., Zitnick, C.L., 2014. Microsoft coco: Common objects in context, in: European Conference on Computer Vision. Springer, pp. 740–755.

Naik, H., Chan, A.H.H., Yang, J., Delacoux, M., Couzin, I.D., Kano, F., Nagy, M., 2023. 3D-POP - An Automated Annotation Approach to Facilitate Markerless 2D-3D Tracking of Freely Moving Birds With Marker-Based Motion Capture. Presented at the Proceedings of the IEEE/CVF Conference on Computer Vision and Pattern Recognition, pp. 21274–21284.

Nakagawa, S., Gillespie, D.O.S., Hatchwell, B.J., Burke, T., 2007. Predictable males and unpredictable females: sex difference in repeatability of parental care in a wild bird population. J. Evol. Biol. 20, 1674–1681.

Schroeder, J., Hsu, Y.-H., Winney, I., Simons, M., Nakagawa, S., Burke, T., 2020. Data from: Predictably philandering females prompt poor paternal provisioning. https://doi.org/10.5061/DRYAD.0T313
